# ExposomeX: Integrative Exposomic Platform Expediates Discovery of “Exposure-Biology-Disease” Nexus

**DOI:** 10.1101/2022.11.23.517586

**Authors:** Bin Wang, Changxin Lan, Guohuan Zhang, Mengyuan Ren, Yanqiu Feng, Ning Gao, Weinan Lin, Bahabake Jiangtulu, Zhijian Liu, Xuqiang Shao, Shu Su, Yuting Wang, Han Wang, Fanrong Zhao, Bo Peng, Xiaotong Ji, Xiaojia Chen, Min Nian, Mingliang Fang

**Author notes:** Correspondence and requests for materials should be addressed to MF.

## Abstract

Exposome has become the hotspot of next-generation health studies. To date, there is no available effective platform to standardize the analysis of exposomic data. In this study, we aim to propose one new framework of exposomic analysis and build up one novel integrated platform “ExposomeX” to expediate the discovery of the “Exposure-Biology-Disease” nexus. We have developed 13 standardized modules to accomplish six major functions including statistical learning (E-STAT), exposome database search (E-DB), mass spectrometry data processing (E-MS), meta-analysis (E-META), biological link via pathway integration and protein-protein interaction (E-BIO) and data visualization (E-VIZ). Using ExposomeX, we can effectively analyze the multiple-dimensional exposomics data and investigate the “Exposure-Biology-Disease” nexus by exploring mediation and interaction effects, understanding statistical and biological mechanisms, strengthening prediction performance, and automatically conducting meta-analysis based on well-established literature databases. The performance of ExposomeX has been well validated by re-analyzing two previous multi-omics studies. Additionally, ExposomeX can efficiently help discover new associations, as well as relevant in-depth biological pathways via protein-protein interaction and gene ontology network analysis. In sum, we have proposed a novel framework for standardized exposomic analysis, which can be accessed using both R and online interactive platform (http://www.exposomex.cn/).

## INTRODUCTION

It has been well acknowledged that environmental factors make enormous contributions to human health. Due to the spatial and temporal differences as well as the highly diverse environmental exposure factors, exposome has become the hotspot of next-generation environmental health study by providing a “perfect” solution to use the entire environmental factors for the relevant human health risk assessment ^1,2^. Exposomic studies usually make use of multiple-dimensional data, including several to hundreds of external exposure factors such as physical, chemical, biological, or even social activities ^3-6^. However, compared to the booming genomics, proteomics, and metabolomics, the characterization and analysis of exposome are still at its infancy and the standardized workflow for exposomic analysis is of high demand.

The roadmap to link thousands of exposure factors with the concerned diseases is the ultimate goal of the exposomic study. However, there are a few major challenges that hurdle its development and application. First, it is of great difficulty in integrating and interpreting the high-dimension data including external exposome, internal exposome, and disease outcomes to build their associations. There are many different types of epidemiological designs for exposomic studies, e.g., cross-sectional study, case-control study, cohort study, longitudinal study, panel study, and time-series study. For these designs, the data pre-treatment and statistical methods are overall distinguished. In addition, different statistical analyses might yield different results, and it is very worth cross-comparing different analytical methods to obtain more robust results with high confidence. Although users can get access to many statistical packages, e.g., Python or R programs, there is still no standardization of statistical method for exposomic studies, which could be very important for the beginners in the field. For thousands of features detected in exposomic analyses, the application of environment-wide association studies (EWAS) assessing exposome–health associations had already shown its capability in terms of the false-discovery proportion and sensitivity ^7^. However, the majority of EWAS analysis is targeted at the association and further in-depth analyses, like mediation, interaction, and prediction, are also very necessary, supporting sufficient data mining and comprehensive utilization of exposomic data. Second, it is challenging to figure out the biological mechanisms of the associations between the external exposure, endogenous change, and the diseases. Besides the statistical associations between exposure factors and diseases, the biological understanding is vital for scientists to further exclude the confounding effect and generate hypotheses for further experimental validation. However, there is no existing platform to effectively integrate and interpret the complex relationships between the exposure, endogenous biological changes, and diseases. During the last decades, there is a big advance in curating the databases of biological targets of xenobiotic chemicals including drugs, toxicants and other chemicals, such as comparative toxicogenomics database (CTD) ^8^ and interaction networks of chemicals and proteins (STITCH) ^9^. Therefore, it is possible to set up this platform to integrate the exposomic with other omics results in a biological way. Third, there is a big demand to efficiently screen the features from mass spectrometry with significant contribution to the concerned diseases. Among all the possible technologies in the exposomic profiling, non-targeted analysis (NTA) has the advantage of high coverage and high throughput and increasingly becomes one major promising tool in exposomoic studies ^10^, though the data analysis is still the bottleneck. Unlike the metabolomics or proteomics analysis, either simple, multiple or pairwise analysis with student-*t* test or *ANOVA* can be conducted to find the dysregulated features among the hundreds to thousands of features. For the exposomic analysis with the complexity of the environmental factors, multiple variants are usually involved, and more advanced statistical methods are needed to prioritize the features that can be linked to the concerned outcomes. Additionally, different statistical analyses may result in distinct results. Therefore, it is very useful to set up multiple statistical analysis and then further conduct the cross-platform comparison. Lastly, it is quite lacking of integrated exposomic databases to expediate the discovery of “Exposure-Biology-Disease” nexus. For example, to assist the completion of the above-mentioned function, a comprehensive database of chemical lists, chemical-protein targets, chemical-disease interaction, and exposure biomarker information are essential. Though those databases can be found in different resources, data cleaning, combination and integration is quite necessary. In addition, the data visualization of different statistical analysis as well as the biological interaction can save lots of time for the data interpretation. Also, it is of utmost need to compare the exposomic result with previous evidences to further confirm or exclude the confounding effect in the correlation analysis using meta-analysis. To date, there is not enough meta-analysis for common environmental chemicals and diseases.

To address those difficulties in the exposomic studies, we aim to standardize exposomic framework and build up one integrated platform “ExposomeX” to bridge the “Exposure-Biology-Disease” nexus (see **Figure 1**). Briefly, it mainly contains six major functions including statistical learning (E-STAT), exposome database search (E-DB), mass spectrometry data processing (E-MS), meta-analysis (E-META), biological link via pathway integration and protein-protein interaction (E-BIO), and data visualization (E-VIZ) **(Figure 1a)**, which are accomplished by 13 analysis modules **(Figure 1b)**. For example, the interaction modes to link the chemical exposure and disease outcome have been demonstrated in **Figure 2**. In sum, “ExposomeX” provides the first most comprehensive analytical platform for the exposomic study including both online web-based user interface and R package **(Figure 1c-d)**.

**Figure 1.**
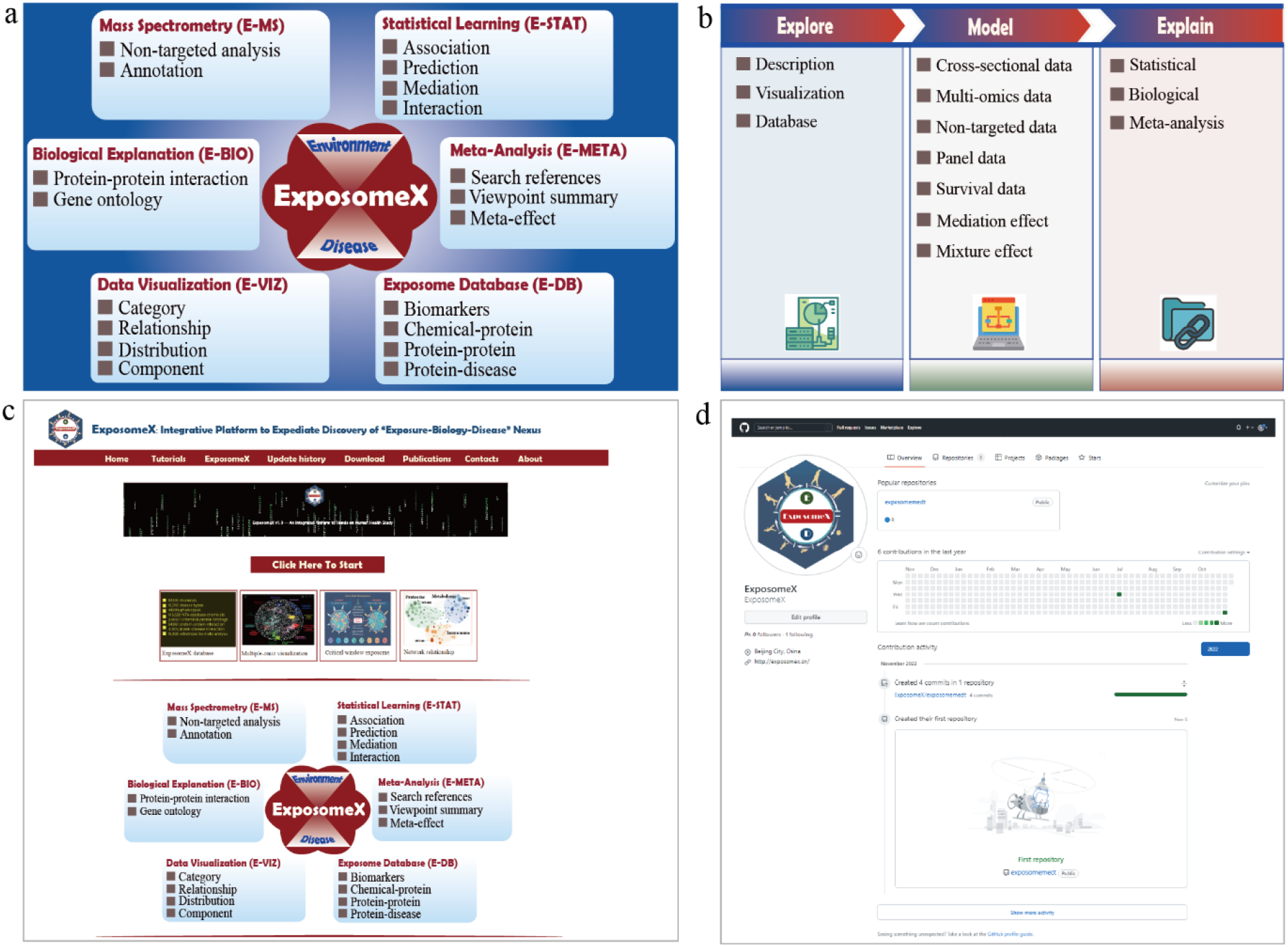
Design of the “ExposomeX” operation platform and its supporting modules. (a) Six functions (i.e., E-MS, E-STAT, E-META, E-DB, E-VIZ, and E-BIO). (b) 13 supporting modules for data exploration (marked as “Explore”), modeling (“Model”), and data interpretation using statistical and biological methods (“Explain”). (c) Web-based user interface (http://www.exposomex.cn/). (d) R package (https://www.github.com/ExposomeX).

**Figure 2.**
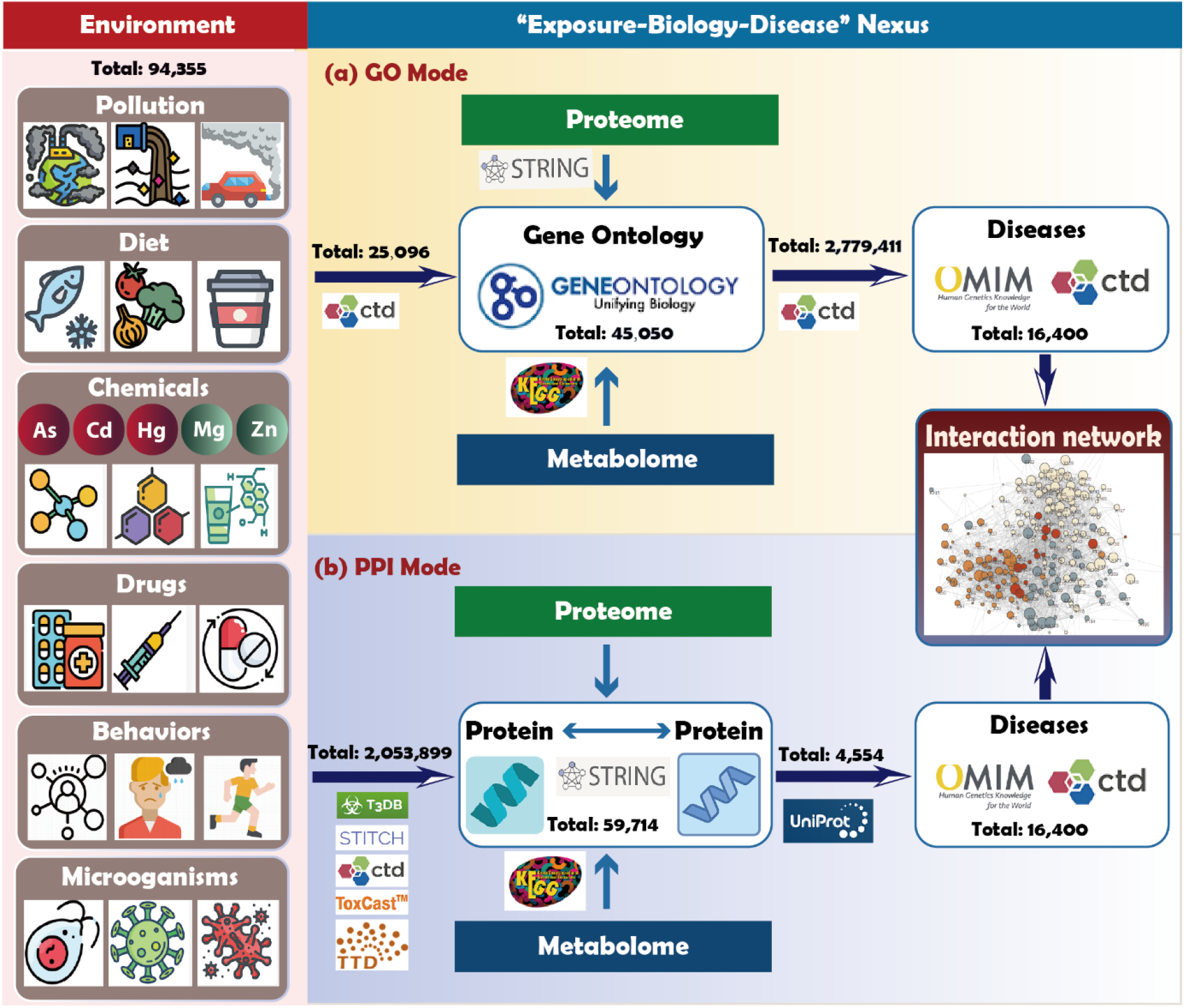
The interaction modes of GO pathways and protein-protein interaction (PPI) to link the chemical exposure and disease outcomes. A total of 94,355 environmental chemicals, 16,400 disease types, 167,654 proteins, and 45,050 GO pathways were included. (a) Gene ontology mode (GO) contains 25,096 chemical-GO pathway interactions. (b) Protein-protein mode (PPI) contains a total of 2,053,899 chemical-protein interactions, 59,714 protein-protein interactions, and 4,554 protein-diseases associations. All the numbers shown in the figure and their sources are accessible in ExpoDB.

## RESULTS

### Curation of exposomic database and its use

To support the development of the six functions of ExposomeX, we have started the framework with curating the databases (ExpoDB) for exposomic chemical search, exposure biomarker database, the targeted proteins of chemicals, chemical-outcomes, and associated proteins of certain diseases. Most of the database are summarized, cleaned, and cross-validated from well-established databases such as CTD ^8^, STITCH ^9^, ToxCast ^13^, and T3DB ^14^. In addition, we have also manually searched the protein targets for other non-chemical environmental exposure; namely, smoking, alcoholic drinking, ozone, NOx, particulate matters, sleeping disorders, heat wave, high-salt diet, exercise, UV irradiation, high-fat diet, and noise. In sum, we have created a total of 2,053,899 pairs of chem-protein interactions from the several major databases and the above-mentioned manual searching, involving 94,355 exposure factors and 167,654 proteins, as detailed in **Table S1 [Supporting Materials (SM)]**. To facilitate the use of ExpoDB, five key sub-functions was further built, including searching the related information (Dictionary), converting the IDs between different databases (Convert), exploring the nexus between exposure, proteins, phenotypes, and diseases (Nexus), annotating the non-targeted features from high-resolution mass spectrometry (Annotation), and learning the nomenclature of “ExposomeX” platform (Abbreviation) (**Figure S1**).

### Exploratory analysis by statistical and visualization modules

As shown in **Figure** 3, we have accomplished the exploratory description analysis of the exposome data under the context of one specific epidemiological design. We started the analysis with the function of “Tidy data”, including data imputation, filling the variables with missing values and manipulating variables with low variance (**Figure 3a**). Following that, data transformation will be conducted for scaling, log transformation, dummy transformation, etc. according to data types and distribution, which can be defined in the data analysis (**Figure 3b**). Subsequently, data can be analyzed for generating Table 1 (summary of population characteristics), extreme value detection, normality test, pairwise or multiple group comparison, and multi-variant correlation using ExpoStatistics and ExpoViz modules (**Figure 3c-d**). Meanwhile, the visualization of those high-dimensional data can be conducted using the simple interactive interface on the webpage. As an example, the data visualization of description, correlation heatmap, network correlation, category, relationship, distribution, and components have been included in the toolkit (**Figure 3e-k**). Most of the common data analysis functions have been incorporated into ExposomeX. The other detailed results are provided in “**Supplementary Data_Fig3A-G**” and “**Supplementary Data_Fig3H-K**”.

**Figure 3.**
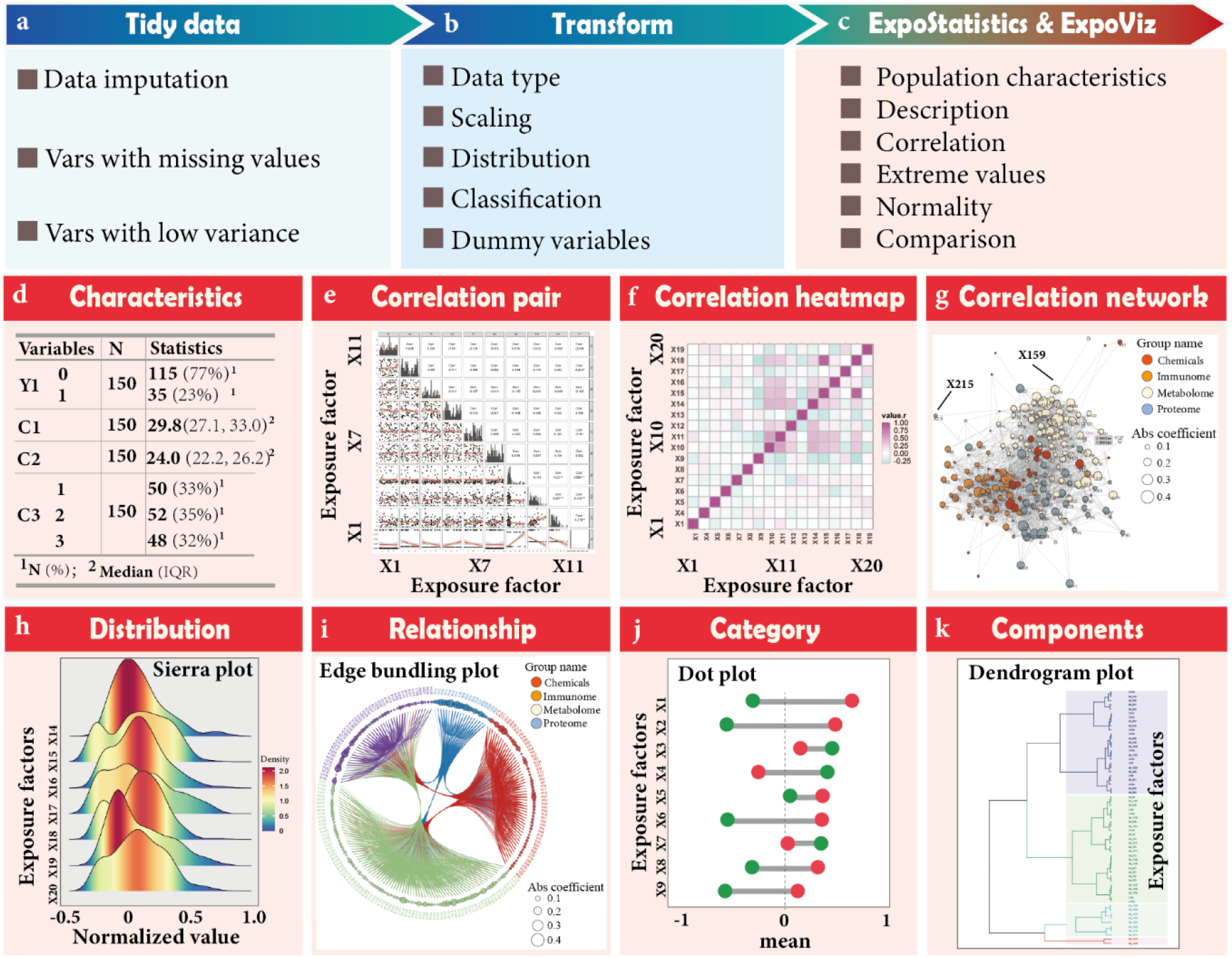
Exploratory analysis by following a specific epidemiological design using ExpoStatistics and ExpoViz modules. (a-c) The basic procedures for cleaning the data for further analysis using the ExpoStatistics module, including tidy data, transform data, and statistical analysis. (d) Standard table of demographic information summary (i.e. Table 1). (e-g) Typical data visualization descriptions using the ExpoViz module, including integrated relationship exploring, correlation heatmap, network relationship between features or between multiple omics. (h-k) The typical visualization of the four classes of visualization including distribution (e.g., sierra plot), relationship (e.g., edge bundling plot), category (e.g., dot plot), and components (e.g., dendrogram plot).

### Standardization of the exposomic analytical framework and workflow

In ExposomeX, we further aim to build the framework and standardize the analytical workflow for various tasks under different epidemiological scenarios (**Figure 4a**). In the model task, we propose to conduct the following functions with the aims of association (e.g., ExpoCros, ExpoSurvival and ExpoPanel), prediction (e.g., ExpoCros, ExpoMultiOmics), multi-omics (i.e., ExpoMultiOmics), non-targeted analysis (i.e., ExpoNTA), mediation (i.e., ExpoMediation), and mixture effect (i.e., ExpoMixEffect), which were determined based on the epidemiological design, data structure, and analytical tasks in the exposomic study. In this stage, ExposomeX mainly contains one specific epidemiological design (cross-sectional, longitudinal, case-control, cohort and case follow-ups), and the data structure (e.g., cross-sectional, panel, and cencored). The statistical learning is supported by the most popular algorithms to obtain the optimal results, including the traditional models (e.g., linear regression, regularization regression, bayesian methods, kernel-based algorithms, association rules learning) and the ensembling models (e.g., decision tree learning, artificial neural networks, ensemble algorithms boosting and bagging, as well as stacked generalization).

**Figure 4.**
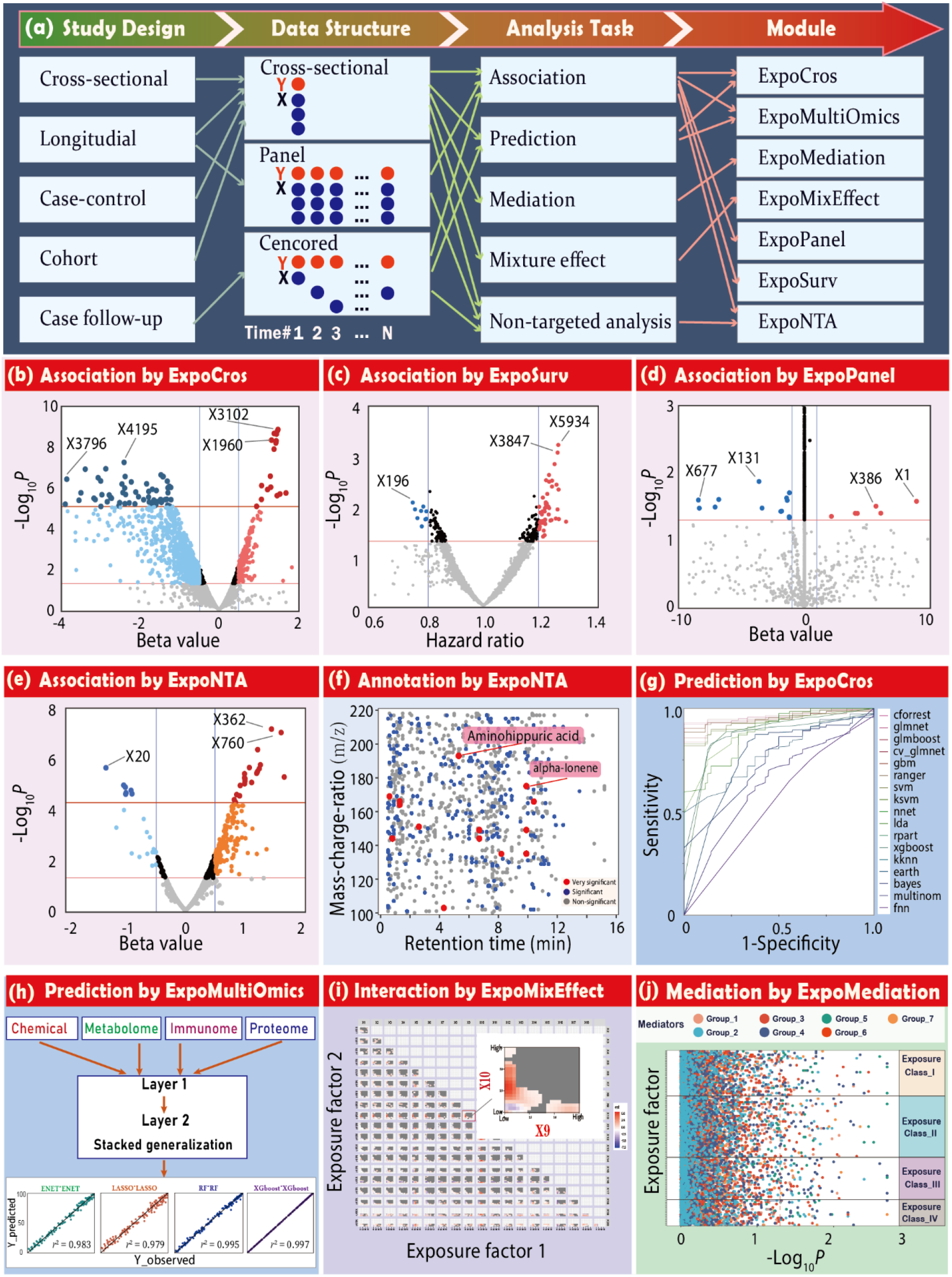
Applying ExposomeX models to solve various tasks and the visualization for some typical results of these modules. (a) The main parameters to choose the appropriate models, mainly include epidemiological study design, data structure, and analytical tasks; (b-e) The association analysis results from the concerned using the four modules of “ExpoCros”, “ExpoSurvival”, “ExpoPanel”, and “ExpoNta”, respectively. (f) Annotating the significant features using the “ExpoNTA” module from the non-target analytical features. (g-h) The prediction analyses for the single-omic and multi-omic data using the modules of “ExpoCros” and “ExpoMultiOmics”, respectively. (i) Exploring the mixture effect between various environmental factors using the “ExpoMixEffect” module. (j) Screening the mediators from the seven potential mediator groups (M1-M7) interacted with the four potential exposure groups (G1-G4) by mediation analysis using the “ExpoMediation” module. For the figures above, “Beta value” stands for the coefficient of linear or Logistic regression model, and “Hazard ratio” for the risk assessment of Cox proportional hazards model.

Hereby, we explain the performance of a few key models. Based on analytical tasks, some typical modeling results are presented in **Figure 4b-j**. The association analysis uses four modules of “ExpoCros”, “ExpoSurvival”, “ExpoPanel”, and “ExpoNTA” **(Figure 4b-e)**. Both positive and negative hits from different types of epidemiological designs can be effectively screened and labeled. In the case of non-targeted analysis, we aimed to develop outcome-directed statistical methods to find the annotated features which warrants in-depth confirmation. As you see in the example (**Figure 4b**), we have correlated about 7,000 features for a cohort study with the health outcome using ExpoCros modules. The result revealed only 58 features can be associated with the health outcome by multiple hypothesis correction, which efficiently prioritize and annotate the massive features in the sample. The prediction analyses for the single-omic and multi-omic data were conducted using the modules of “ExpoCros” and “ExpoMultiOmics”; respectively **(Figure 4 f-g)**. In the case of ExpoCros, ROC curve from multiple models with different sensitivities and specificities can be readily displayed in the ExposomeX. For the multiple omics, stack generalization (SG) models were used as one example to integrate and compare the output from different sub-models for the exposure groups of chemicals, metabolome, proteome, and immunome. For the multi-omics study, the high dimensionality of data is a problem that cannot be ignored. We provided two strategies for dimensionality reduction including regularized variable selection and variable importance screening. Intra-omic and inter-omics network were provided to find potential biological associations within each single exposure group and between different exposure groups. The results are more likely to be reliable, unbiased, and reproducible. For the mixture model, we have included several most commonly used methods such as stepwise multiple-factor linear regression (MLR), least absolute shrinkage and selection operator (LASSO), elastic net (ENET), weighted quantile sum (WQS) regression, and Bayesian kernel machine regression (BKMR), which enable users to apply different models for exploring their mixture effect, as well as the interaction effects between concerned features on the target outcome. For instance, the heatmap shows the interaction effect between X1 and X2 within all concentration ranges marked with different colors (**Figure 4i**). For the mediation effect, there is a critical knowledge gap for addressing a multivariate mediation setting where there are not only high dimensional mediators, but also multiple toxicants with inherent collinearity. Our study aims to advance methodological applications to investigate a mixtures mediation setting with multiple exposure classes and groups of endogenous biomarker mediators. We make use of an advanced analytical framework that integrates environmental risk score construction with the aforementioned multivariate mediation analysis methods **(Figure 4j)**. In this framework, we provide a guided discussion on approaches that include one-at-a-time pairwise mediation, exposure dimension reduction, and mediator shrinkage, dimension reduction, and penalization. The other detailed results are provided in “**Supplementary Data_Fig4B**” to “**Supplementary Data_Fig4J**”.

### Comprehensive explanation analysis of “Exposure-Biology-Diseases” Nexus

Statistical analysis was conducted to determine the contributions of the environmental factors and their partial dose-response relationship with the target outcome **(Figure 5a-c)**. In addition to the statistical analyses, we further would like to explain the statistical result based on previous epidemiological studies using our “MetaRefer” and “MetaReview” (e.g., zinc intake and risk of asthma) functions (**Figure 5d-e**). Likewise, in the “ExpoMeta” module, we also set up a meta-analysis “MetaEffect” for the association between copper exposure and the risk of polycystic ovary syndrome by including 12 studies. The individual results were shown in the forest plot and summarized using both fixed-effect and random-effect models considering their contribution weight **(Figure 5f)**. Due to the meta-analyses and potential new studies in the future, we also use the interactive mode to allow the users to upload some new studies in the system and integrate the information by sending us the information according to the standardized forms (http://www.exposomex.cn/#/contact).

**Figure 5.**
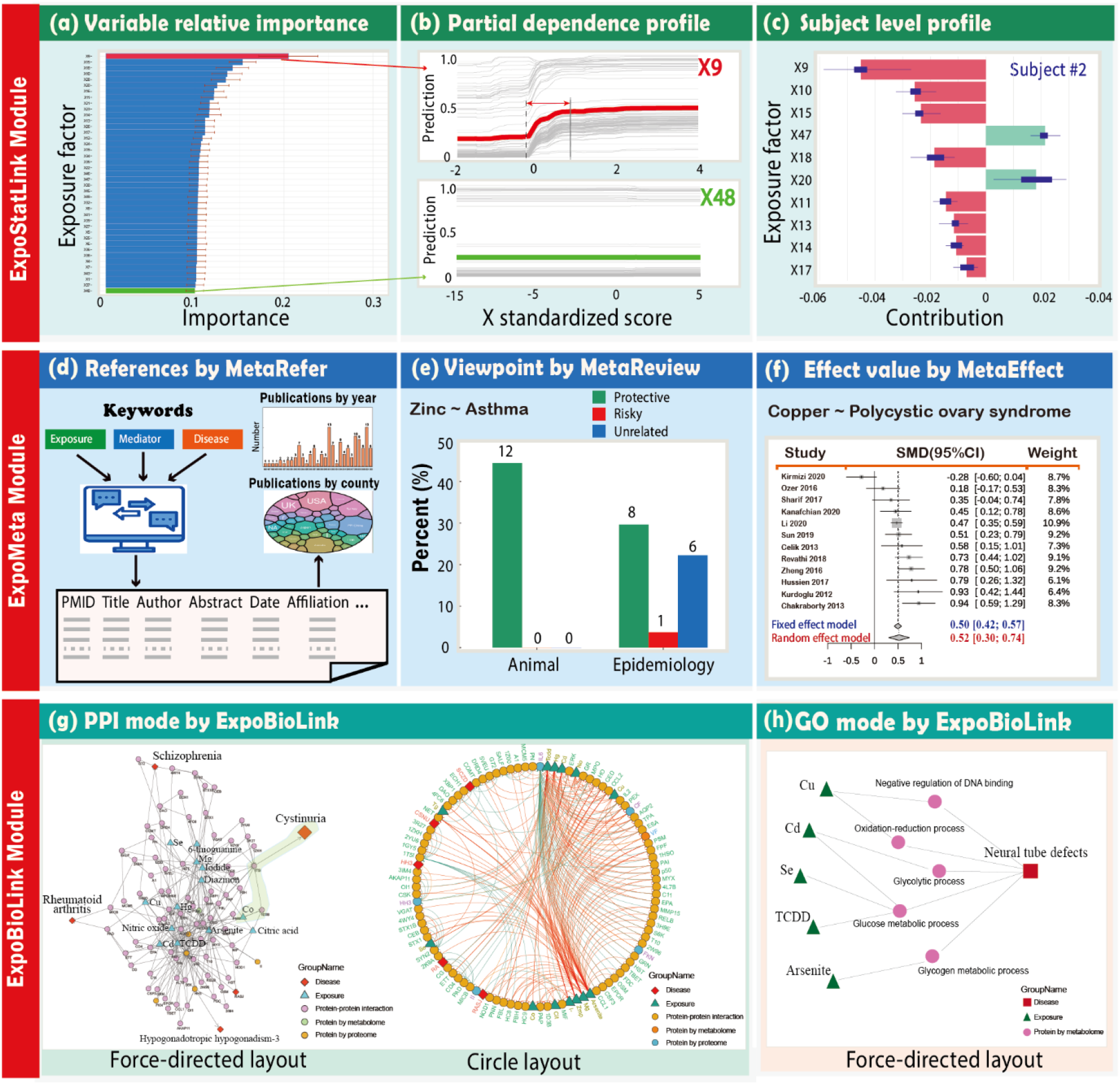
Explain model results by statistical analysis, meta-analysis and biological ways. (a) Statistical analysis to determine the relative importance of the environmental factors. (b) The partial dependence profile of two factors. (c) Contributions of the concerned variables to the outcome, e.g., Subject #2. (d-f) Auto-search, review and meta-analysis of the previous publications based on the data-mining technology and the established literature database. (g-h) Biological explanations of the multi-omics data using the protein-protein interaction (PPI) and gene ontology (GO) modes with the force-directed or circle layout for environmental exposure chemicals and various disease outcomes. The terms starting with “X” (e.g., X9 and X48) represent exposure factors as detailed in **Supplementary Data_Fig5GH**.

GO and PPI networks were used as the two major methods to integrate the “Exposure-Biology-Diseases” nexus. As an example, we have investigated the biological network understanding of 13 environmental factors and six diseases as shown in force-directed and circle layouts (**Figure 5g**). Here, a couple of environmental chemicals can work on the same or relevant protein targets and be linked with the diseases, suggesting the mode of action for the chemical exposure and possible chemical-chemical interaction. In the second example, we used the GO network analysis and found the possible mechanism for the toxic metal induced neural tube defect (NTD) development. All of those toxic metals have been associated with NTD in previous epidemiological studies and the disrupted protein/metabolic pathways as shown in **Figure 5h**, suggesting possible links between those environmental factors and the diseases. Overall, this platform can provide a useful mapping for the possible mechanism of chemical-disease network. The other detailed results were provided in “**Supplementary Data_Fig 5ABC**” to “**Supplementary Data_Fig 5GH**”.

### Performance validation of ExposomeX for multiple-dimensional dataset

To further validate the performance of ExposomeX in the processing of multiple-dimensional datasets, we have re-analyzed two previous studies with typical epidemiological designs and multiple-omics data. First, we conducted a mediation analysis using the “ExpoMediation” module following the study ^11^, which includes various mediation analysis methods by reducing the dimensions of exposures and mediators. The data in this previous study were adopted, including a total of 38 environmental toxicants (phthalates, phenols and parabens, polycyclic aromatic hydrocarbons, and trace metals) and 61 mediators (cyclooxygenase pathway, cytochrome P450 pathway, lipoxygenase pathway, parent lipid compounds, and inflammatory biomarkers). The analytical results are shown in **Figure 6**, which can reproduce the data of Figure 3, Figure 5, and Table 3 in Aung et al. (2020)^11^. In **Figure 6a**, the significant mediation pathways were screened. After dimension reductions for both exposure and mediation variables, the overall mediation effect between these groups was summarized in **Figure 6b**, and the detailed mediators were shown in **Figure** 6c. In addition, we further conducted a few more in-depth analyses based on the mediation result. For example, **Figure 6d** showed the interaction between the environmental factors and the results suggested 2-OH-FUE and 2/3-OH-PHE might synergistically work on the outcome of congestion weeks at low concentrations, while there is no significant effect at other ranges. To further illustrate how these environmental factors can be biologically linked with the early labor, we have pulled out the proteins that have been reported to be linked with the reduced congestion weeks and conducted the PPI network analysis. As shown in **Figure 6e**, four proteins and three pathways might be one possible mode of action to establish the “exposure-biology-diseases” nexus for the two exposure factors manganese (Mn) and arsenic (As). **Figure 6f** further quantified the contribution of nine phthalates to the gestational week, and the negative partial-dependence relationship between two typical phthalates and gestational week was shown. Finally, the finding of the association between a typical phthalate of mono-n-butyl phthalate and risk of preterm birth in previous studies has been summarized using meta-analysis and the forest tree with the overall odd ratios of 1.22 (95%CI: 1.01-1.49), indicating their positive association of the previous studies. The other detailed results were provided in “**Supplementary Data_Fig6ABC**” to “**Supplementary Data_Fig6G**”.

**Figure 6.**
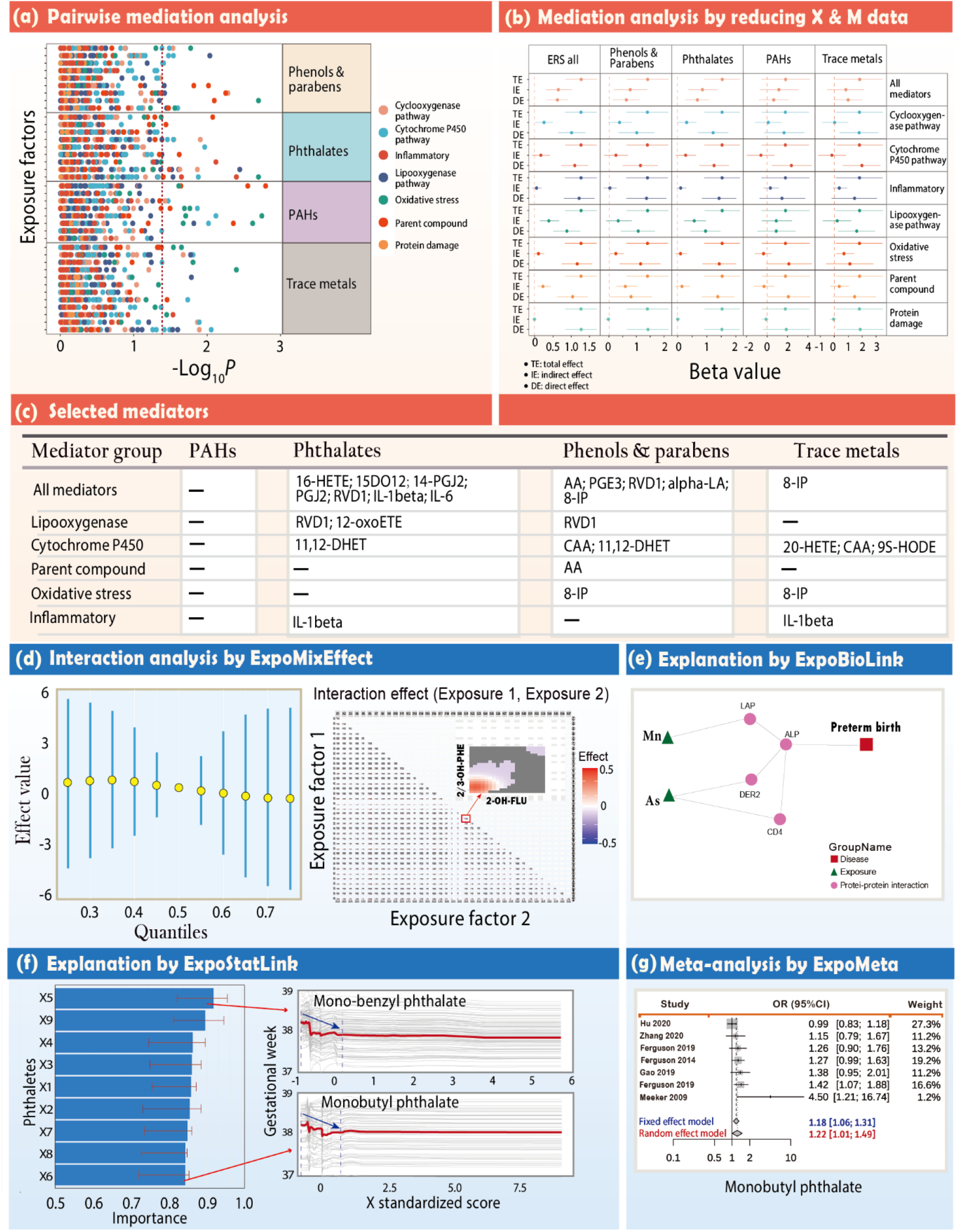
Applying ExposomeX platform to repeat the mediation analysis and conduct new in-depth analysis for the data from the published study ^11^. (a) Estimated -log_10_(*P*-values) of natural indirect effects from pairwise weighted mediation models for gestational age at delivery. (b) Forest plot of mediation analysis for environmental risk scores, mediator group effects, and gestational age at delivery. (c) Pairs of environmental risk scores and mediators selected by joint significance test in “*hima*” package. (d) Interactions analysis between various exposure factors to gestational age at delivery using the module of “ExpoMixEffect”. Metabolites of fluorene and phenanthrene (i.e, 2-OH-FLU and 2/3-OH-PHE) were highlighted as one pair of examples. (e) Biological pathway by PPI mode analysis using the module of “ExpoBiolink” to understand possible mechanism of chemical exposure, endogenous change, and pre-term birth”. (f) Statistical explanation analysis using the module of “ExpoStatLink” to explain the variance importance for the nine concerned phthalates, and the partial dose-response dependence between two phthalates and gestational weeks were provided, of which the sensitive concentration range was marked by blue arrow. (g) Forest plot of OR (95% CI) using the module of “ExpoMeta” on the mono-n-butyl phthalate on the risk of preterm birth.

Second, we validated the overall functions of multiple-omics analysis, association, feature prioritization of NTA, statistical explanation, prediction, and mixture effect deconvolution using multiple relevant modules. The data from the previous study (noted as “MO-Study”) ^12^ including metabolome, proteome, and immunome information was adopted. In **Figure** 7, we present the typical results using SG models, including LASSO, ENET, Random Forest, and Xgboost. Their model evaluation of *R*^2^ by the 5-fold cross-validation for the four models were provided, in **Figure 7a**, respectively. We can see that the values of *R*^2^ for the four models were above 0.98, and even up to 0.999 for XGboost, which is overall better than that in **Figure** 3B of the original MO-Study. As we can see that there are high correlations between each other in the inter-omic features (see **Figure** 4 in MO-Study), some other features with high correlations with TL may be chosen when using a new algorithm. In **Figure 7b**, the network relationships between the important features selected by the SG model (XGboost * XGboost) were depicted. Metabolomic and proteomic features had closer relationships compared to those with immunomic features. From the correlation matrix of all features, we filtered different thresholds to visualize the connections between the different omics. For XGboost model, the scatter plot of variable importance in the model and the *P* value of the correlation between the individual features and time to labor (TL) were shown in **Figure 7c** (left panel), and the partial-dependence relationships between three features and TL were shown in **Figure 7c** (right panel). We found that the selected top important variables for prediction using XGboost model are dramatically different from those in MO-Study ^12^. The models of mediation and mixture effects were further conducted using the analysis modules of “ExpoMediation” and “ExpoMixEffect”, respectively (**Figure 7d-e**). For the prioritization of NTA features (**Figure 7f**), we are also able to find more significant features associated with TL after applying various statistical models, compared with the results in MO-Study. In sum, our platform can efficiently conduct the multiple-omic analysis with various models and explore more underlying information. The other detailed results were provided in “**Supplementary Data_Fig7A**” to “**Supplementary Data_Fig7F**”.

**Figure 7.**
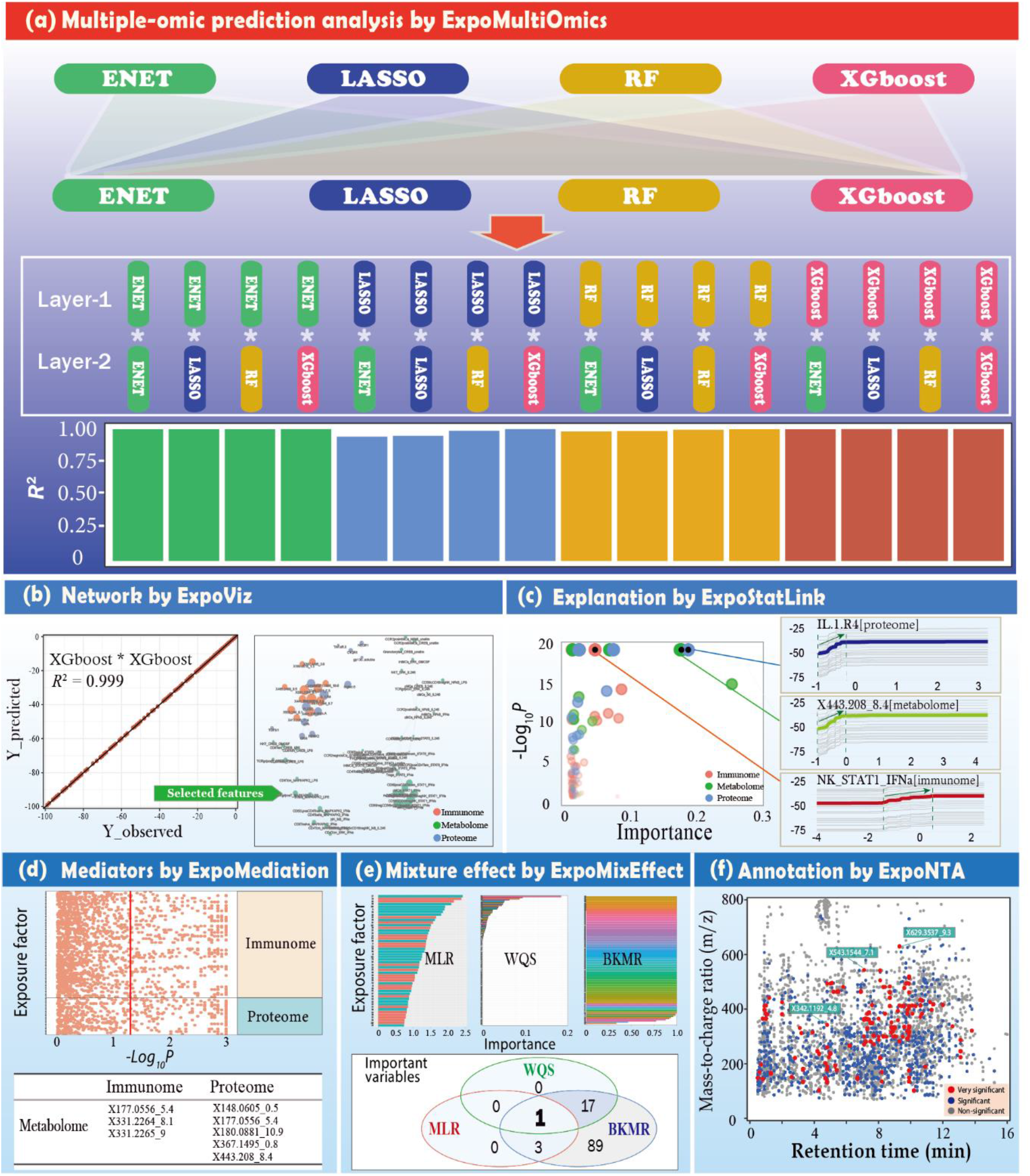
Applying ExposomeX platform to re-analyze and conduct new analysis for the data from the published study ^12^. (a) Comparison of the prediction performances of the 16 SG integrated models based on the combination of LASSO, ENET, RF and XGboost; (b) Network analysis for the selected features from three omics of immunome, metabolome, and metabolome analyzed by XGboost*XGboost and demonstration with ExpoViz module; (c) Statistical explanation of the selected features for the final prediction model (SG: XGboost*XGboost); (d) Pairwise mediation analysis between the significant features of proteome, immunome and exposure factors; (e) The importance of the proteins contributing to the time to labor (TL) from the three models of MLR, WQS, and BKMR, as well as their interaction effect on the TL by the module of “ExpoMixEffect”. (f) Scatter plot of the mass-charge-ratio and the retention time with the significant HRMS features for the metabolomic data annotated by the module of “ExpoNTA”.

It should be also noted that the main statistical platform in our study was developed based on the foremost framework of R package “mlr3”, which embraces the most advanced algorithms to facilitate the modeling robustness and accuracy. Most of the analysis can be conducted very efficiently and the re-analysis of both demonstration data can be accomplished within several minutes, indicating the high performance of ExposomeX in processing the multi-dimensional data.

## DISCUSSIONS

Exposome has become a hot topic since it was coined in 2005. Though still controversial, it has been well acknowledged that environmental factors can explain largely for the disease burden and exposome has been regarded as one ultimate solution to decipher the real contributions of environmental factors to diseases. However, to date, there are very few successful applications of exposome in practice and most of the publications are either perspectives or review articles. The exposomic study design and the subsequent data analysis of multiple dimension data seems to be more difficult than its counterpart omics such as metabolomics and proteomics. There are a couple of major challenges for exposome study. As mentioned above, those primarily include the integration and interpretation of high-dimension data, understanding the biological mechanisms of the associations between the external exposure, endogenous change, and the diseases, identification of the features from NTA with significant contribution to the concerned diseases, and the integration of all the associated databases and visualization tools. It should be also noted that the exposome is a very interdisciplinary field which involves the knowledge from epidemiologists, clinicians, statisticians, chemists, geologists, economist, sociologists, and toxicologists. It is very challenging for one scientist to acquire all the needed skills. Researchers who currently have exposure omics research demand do not have this systematic statistical background, and are prone to misuse relevant models, which in turn affects the accuracy of the results. In addition, different statistical analyses can result in various outputs and single or biased analytical methods are likely to lead to the loss of useful information in the exposomic data. With the developed platform ExposomeX targeting at those challenges, the boundary and bottleneck of the exposome for the scientists from different fields have been greatly improved.

In this study, we have conceptualized the exposome analytical workflow and built the most integrated platform “ExposomeX” to connect “Exposure-Biology-Disease” nexus. Specifically, E-STAT aims to correlate the exposomic factors with the diseases using different layers of statistical analyses with a total of over 100 different statistical models for epidemiological studies of cohort, case-control, and cross-section to complete the tasks of association, mediation and prediction. E-DB provides the function to search the exposure chemicals (∼ 100,000) by mass-charge-ratio, exposure biomarkers (∼ 1,500), protein targets and protein-protein interaction (∼ 60,000) curated from previous databases and literature search. E-MS can prioritize and annotate the features that are correlated with the diseases under different epidemiological design from the large exposomics data acquired by high resolution mass spectrometry. E-META serves as a tool for the meta-analysis on the result from previous epidemiological results with ∼ 20,000 research articles by now. E-BIO provides the tool to integrate the exposomic and other omics result via pathway integration and protein-protein interaction. E-VIZ enables the visualization of the statistical and biological analysis. Using our platform, the users can complete almost all the key functions of exposomic studies, including mediation, interaction, and prediction analyses of multiple-dimension data, and biological understanding for the statistical correlation. For the epidemiological design, the most common designs including cross-sectional, longitudinal, case-control, cohort and case follow-ups are included. Its excellent performance has been validated in the re-analysis of two previous studies. In both cases, besides the fast completion of the tasks as required by the original studies within a few minutes, ExposomeX was able to provide another level of understanding and confidence level of the association found in the studies with additional newly mined information.

ExposomeX does not aim to “reinvent the wheel” and is novel in a few aspects. First of all, we have set up the first systematic exposomic framework for statistical analyses and biological analyses, which enables the users to effectively mine the useful information hidden in the exposomic data and tentatively establish the “exposure-biology-diseases” nexus. The modules and their specific adopted methods are based on the existing knowledge on the epidemiological studies and data structures of a typical exposomic study. To facilitate the biological understanding, we finally decide to use GO pathway enrichment and PPI network to link the chemicals, biological pathways, and diseases due to the availability of those data from previous studies, such as the information on the transcriptomics data and binding or regulatory proteins of chemicals or other environmental factors. This network analysis has not been well recognized in the exposomic studies, which mainly focus on the association between different omics results. Second, ExposomeX can provide the re-analysis of various statistical models and conduct the multifaceted task analysis by benchmarking their performance using the same resampling dataset, which enables the users to find more useful and novel information in an unbiased manner. Third, ExposomeX also provides the largest database fusion for exposomics studies by curating, cleaning and integrating the existing database as well as mining the new information. Besides the existing database from other databases such as CTD, ToxCast and STITCH, we have collated more information from literature search under the exposomics concept such as PM_2.5_, sleeping disorders and radiation, which are all key exposomic factors. Likewise, meta-analysis is another novel database we have created in the ExposomeX. Via this function, we are able to reuse the value of previous thousands of environmental epidemiological studies that have been conducted. We also allow the user to upload their meta-analysis result, which can further expand the meta function in the ExposomeX. Last but not the least, all the modules developed in ExposomeX are integrated together for the first time and output of each module can be used as the output of other relevant one, which greatly expediates the analytical workflow and makes it more straightforward to analyze and reproducible using a code script.

The online web form of ExposomeX is very user-friendly and open access. Both R and Web based format of Exposome has been implemented and this standardization greatly reduces the technical bar of exposome by providing more extendable, repeatable, and stable result. This will benefit the majority of these investigators without the bioinformatic expertise that has been required to process exposomic data by using command-line driven software programs. The novel online platform to process exposomic data uses an intuitive graphical interface and does not require installation or technical expertise. This platform allows users to easily upload and process the epidemiological data, liquid chromatography/mass spectrometry, disease outcome data with only a few mouse clicks. For each input, one example data and detailed standard operation procedures have been provided. The users can easily formulate their information to the required format and complete the process. Results can be browsed online in an interactive, customizable table showing statistics, putative compound identities for each feature, omics integration and “exposure-biology-disease” network. Additionally, all results and images can be downloaded as zip files for offline analysis and publication. The platform can supply good resources for both advanced and beginner users. In addition, the job can also be shared between different users. Comparing with other omics platforms such as XCMS and Metaboanalyst with thousands of users during last several years, it can be expected that ExposomeX will attract lots of users.

In summary, the ExposomeX is an ecosystem platform of R packages and online web interaction that share an underlying design philosophy, grammar, and uniform data structure, which provides a comprehensive, transparent, traceable, reproducible, and object-oriented computational framework for exposomic data processing and analysis. ExposomeX can provide great benefit for the exposomic field, particularly in the analytical framework standardization, data sharing, and reproducible analyses. In addition, the flexibility and extensibility are another focus of ExposomeX. The design concept allows for the easy integration of other tools with ExposomeX, therefore making ExposomeX flexible and extensible within the exposomic community. In our future version of our platform, we are going to further increase our capability in the exposomics studies by updating the statistical methods and all different kinds of databases we have applied, especially the epigenetics modulation by the environmental factors. Also, literature database of meta-analysis can be enlarged with involvement of users. With those improvements, we are likely to establish a platform to interpret all the exposmic studies and bring the exposomic study in practice.

## METHODS

### Exposome Database (E-DB)

To support the development of the six functions of ExposomeX, we firstly curated the databases for exposomic chemical search, exposure biomarker database, the targeted proteins of chemicals, chemical-outcomes, and associated proteins of certain diseases. Due to the large number of chemicals curated from different databases, we have assigned one unique ID, e.g. EXE00001-EXE94355 for exposure factors, as shown in the supplementary data file “DB_Exposure_Unfold.csv”. All the database files can be downloaded from the website: http://www.exposomex.cn/#/download.

### Exposomic chemical database

The chemical databases are mostly derived from our previous study on the HExPMDB ^15^. Briefly, we screened as many chemicals as possible in different databases and sources relevant to exposome research including environmental pollutants, toxicants, gut microbiome-derived metabolites, disinfection by-products, carcinogens, food nutrition and additives. Most of these compounds are chemicals of concerns due to either their production volume or toxicity. A total of 94,355 unique chemicals were obtained, among which 68,220 chemicals with Chemical Abstracts Service Registry Number (CASRN) were included. Additional information includes chemical identifiers (DTXSID, chemical name, CASRN, InChIKey and IUPAC name), structures (Simplified Molecular Input Line Entry Specification (SMILES) and InChI string) and intrinsic properties (molecular formula, average mass and monoisotopic mass).

### Biomarker database

Exposure biomarker database is from our previous study ^16^. Briefly, we established the exogenous metabolism of more than 1,600 compounds after two rounds of literature searches and data fusion of existing public databases. Among them, the first round of literature search aimed at 385 typical environmental pollutants, and collected 3,475 literatures related to exogenous metabolism of environmental pollutants; the second round of literature search used six keywords (such as “metabolism”, “metabolites”, “Biotransformation”, “Microsomes”, “Toxokinetics”, and “Pharmacokinetics”) together with compounds on the U.S. EPA high production volume (HPV) list, with a collection of over 400,000 articles on the PubMed website. Compound search, similarity search and prediction function are also available in the provided link http://www.ecobiotrans.asia/, which was hyperlinked in the ExposomeX.

### Chemical-protein targets

Since the protein targets of environmental factors are the key components for the protein-protein interaction (PPI) network. To do this, we first curated and cleaned all the chemical protein targets from several previous major databases, such as T3DB ^14^, STITCH ^9^, CTD ^8^, ToxCast databases ^13^, and drug targets ^17^. Both target proteins and regulating proteins have been collected and categorized in the database. In addition, we have manually searched for new protein targets using keywords such as “protein targets”, “chemical proteomics” and others from previous studies. Besides, we have also searched the protein targets for other non-chemical environmental exposure; namely, smoking, drinking, ozone, NOx, PM, sleeping disorders, heat wave, high-salt diet, exercise, UV irradiation, high-fat diet, COVID-19, and noise. All of those factors have been known to be very important factors in exposome. For them, a total of 70,073 pairs of chem-protein interactions were adopted for the association database by our manual confirmation. In this version, we have summarized a total of 2,053,899 pairs of chem-protein interactions from the several major databases and the above-mentioned manual searching, involving 94,355 exposure factors and 167,654 proteins, as detailed in **Table S1**. Binding protein and regulating proteins are separately marked in the PPI network.

### Chemical-outcome

In this platform, we have collected the information of common diseases and non-diseased phenotypes in the exposomic analysis. Due to the large number of diseases for human, we only selected those who have been known as the ones that can be correlated with the environmental exposure. The diseases list a total of 16,400 subtypes of diseases covering almost all the disease types with close relationship with environmental exposures. The disease names follow the OMIM standards (http://www.ncbi.nlm.nih.gov/omim/), or MESH ID (https://meshb.nlm.nih.gov/) referring to CTD database ^8^ and their IDs were inherited. Also, we added some important outcomes of wide concern but without clear OMIM or MESH ID, e.g., spontaneous preterm birth, and decreased gestational week. We included generally non-lethal diseases, which includes but not limited to non-communicable diseases, reproductive and childbirth diseases (e.g., infertility; early puberty, preterm birth, low birth weight), mental disorders, autoimmune diseases, and obesity. For the chemical-outcome association, part of them were inherited from chemical-disease information of the CTD database, but additional association information was further searched from the original paper. **Protein-outcome relationship**. To include the diseases related proteins which can be used for further protein-protein interaction and pathway analysis, we collected the protein targets of diseases as indicated above by curating their targets from UniProtKB ^18^ and NCBI proteins (https://www.ncbi.nlm.nih.gov/). Till now, we adopted a total of 4,554 protein-disease associations, including 4,446 proteins and 4,220 diseases. The user can further increase the disease types and associated proteins if it is not included in the database. For example, we added 75 associations between protein and spontaneous preterm birth to better understand the relationships in **Fig 6e**.

### Mass Spectrometry Data Analysis (E-MS)

#### Mass spectrometry search and compound annotation

We have enabled the users to search compounds in CASRN, formula, mass-charge-ratio (m/z), adduct search, and accuracy (in ppm). A total of 31 adducts for the positive mode (e.g., M+H, M+3H, M+H+2Na, and M+ACN+2H) and 15 adducts for the negative mode (e.g., M-3H, M-2H, M+Na-2H, and M+FA-H) are considered to infer the possible exact molecular weights of the molecular ions, which will be used for peak annotations.

#### Outcome directed non-targeted analysis (NTA)

In this part, we aimed to develop outcome directed statistical methods to prioritize the m/z features that can be linked with the disease outcome. Firstly, the peak picking and alignment can be conducted using the “*XCMS*” R packages and the output table with alignment features was uploaded onto our system. In addition, all the variants including the genders, behaviours and others were all included in the model. The parameters of diseases were included as the output. According to the exposomic study design, different statistical analyses including linear regression model and more sophisticated machine learning model have been all included. For single-factor method, generalized linear regression is used to obtain the association between each factor and outcome, which are further screened by multiple-hypothesis test correction. For multiple-factor method, stepwise, LASSO, and random forest are adopted to screen the features with high associations with the outcome. In summary, for each analysis model, the feature can be statistically correlated with the diseases will be highlighted and compared. Finally, the highlighted features will be further searched against the above-mentioned mass spectrometry database search. Due to the fact that either exogenous or endogenous compounds can exist in the NTA analysis, both databases (HExpMetDB ^15^ and HMDB Database ^19^) have been included in the analysis.

#### Statistical Analysis (E-STAT)

From the perspective of the statistical model, there are five concerned typical analysis tasks including association, prediction, mediation, mixture effect, and explanation, which almost cover all the analysis requirement of exposome study. For the traditional environmental epidemiological study, the key task is to build the association between environmental exposure and target outcomes for the health risk assessment. In our platform, the association can be obtained by the modules of “ExpoCros”, “ExpoPanel”, and “ExpoSurvival” depending on the study design and data structure. With the fast development of the exposomics, massive information has been added into the exposure dataset. It is possible to predict the target outcome due to the large contribution of environmental factors to various diseases. In the modules of “ExpoCros”, “ExpoSurvival”, and “ExpoMultiOmics”, various prediction models by integrating the appropriate machine learning algorithms. For the exposomic study, the underlying mechanism can be explored from the high dimensional data. For example, the effects of the mediation pathway and chemical-chemical interaction on the health outcomes are two important perspectives to understand the potential toxic metabolism. For these demands, the modules of “ExpoMediation” and “ExpoMixEffect” can conduct the mediation and mixture effect analyses, respectively. For the partial dependence of the individual environmental factors, it is useful to know its sensitive exposure range associated with the health outcome, as well as the relative contributions, which can be implemented by the module of “ExpoStatLink”.

#### Meta-analysis (E-META)

It is of utmost need to compare the exposomic result with the previous relevant evidences to further confirm or tell the confounding effect in the association analysis. According to these requirements, the “ExpoMeta” module provide three functions: (1) MetaRef: Auto-search all the relative publications from the important database or journals, e.g., PubMed, Web of Sciences and some other specified prestigious journals based on the input keywords, which has been summarized by the machine learning method and further conveyed to the users. (2) MetaReview: We have been building a standardized database to summarize the up-to-now knowledge about the relationship between environmental exposure and specific diseases. For each topic, the viewpoints from the animal and epidemiological studies are comprehensively summarized by well-trained researchers. (3) MetaEffect: For some issues, if their epidemiological studies have been well-conducted, e.g., association between PM_2.5_ exposure and mortality, the meta-analysis can be conducted to summarize the effect value [e.g., odds ratio (OR), relative risk (RR), hazardous risk (HR)]. The related publications are also summarized in the same database with the function MetaReview. In total, we have summarized 19,781 publications of research articles, of which a total of 449 publications were added by our manual search. All of those publications were downloaded and the main information (e.g., effect values and study years) were extracted and double-confirmed if available.. We also provided an interactive interface for the users to upload their findings by following our input template and the meta-analysis will be automatically updated.

#### Biological explanation (E-BIO)

In this module, we proposed to utilize pathway enrichment of omics result and protein-protein interaction network to link the environmental exposure and disease outcome. The overall outline was displayed in **Figure** 2.

#### GO analysis

Pathway enrichment analysis was used to figure out the GO pathways of each omics data. For gene/proteins, StringDB database was used as the background for pathway enrichment. For the metabolites, KEGG pathways and their corresponding metabolites adopted for enrichment analysis. For the common disease and non-diseased phenotypes, the associated GO pathways have been downloaded from STITCH databased. For the GO of environmental chemicals, its GO pathways have been adopted from the CTD databases. Fisher’s Exact Test was used as the algorithm for the pathway enrichment. The GO pathways from exposure chemicals, different omics results, and diseases were compared, and the matched pathways were highlighted and returned into the output table. In addition to the unsupervised GO analysis, a supervised approach was also adopted by mapping genes/proteins into some cellular toxicity pathways with the end points including adaptive stress response, immune response, and endocrine disruption due to the fact that the traditional gene set enrichment analysis (GESA) might not be able to capture the mode of action of environmental chemicals.

#### PPI network

We further proposed to use PPI as the major tool to integrate the exposomics, metabolomics, proteomics, and target diseases together. The major function was completed using StringDB as an inserted R packages in our program. Basically, the protein targets of environmental exposure were collected from the ExpoDB. For the metabolome, the relevant enzymes of the enrichment pathways were acquired from KEGG enzyme database and input as the proteins involved in the metabolic pathways. Together with genes/protein and diseases associated proteins, the PPI network was calculated using the PPI database. The result can be downloaded as an independent file. Due to the experimental design and availability of some omics data, a few scenarios have been assumed in this platform: (1) There are only exposure factors + diseases; (2) exposure factors + metabolome + diseases; (3) exposure factors + proteome (transcriptome) + diseases; and (4) exposure factors + proteome (transcriptome) + metabolome + diseases. The output result has the number of matched genes for each metabolic pathway. These matched genes can be sent to the subsequent PPI data input.

#### Visualization (E-VIZ)

The visualization of the high dimension data is also very useful for the users in the field. The data visualization of different statistical analyses as well as the biological interaction can save lots of time for the data interpretation. Here, we mainly classify all the visualization task into four types, including category (distinguishing the characteristics of various groups), relationship (statistical relationship between various features), distribution (the distribution characters of the single or multiple factors), and components (the inclusion relationship between the part components and the whole). For each type, users only need to provide the original dataset by following the template and the target types, the “ExpoViz” module can generate various potential visualization types.

#### Ecosystem of “ExposomeX” R program and Online “ExposomeX”

The backend R program is designed by embracing the encapsulated object-oriented programming of R6 classes, data manipulation edged tools of “*data*.*table*” and “*tidyverse*” packages, and advanced machine learning framework of “*mlr3*” package. For “*mlr3*”, it can intelligently select and tune the most appropriate technique for a task, and perform large-scale comparisons that enable meta-learning. Its advanced functionality includes hyperparameter tuning and feature selections, as well as the natively supported parallelization of many operations. Therefore, our platform can provide object-oriented, and extensible framework for various tasks of classification, regression, and survival analysis. The return returned results of R6 classes to users can be further processed by the commonly used R packages with high expansibility and reproducibility.

All the six functions mentioned above written in R program were incorporated into the web-based server to facilitate the use of ExposomeX. The web interface was developed using Vue.js technology (https://cn.vuejs.org/). The backend statistical computing and visualization operations were carried out using functions from the R packages. The integration between browser and web server was established through Python, and the web server and R backend were linked by “*RestRserve*” package (**Figure** 1).

## Supporting information

Tutoriols

## Data availability

All the data for how to use ExposomeX web modules can be accessible on the ExposomeX website: http://www.exposomex.cn/#/download, and the analysis modules can be used at the website: http://www.exposomex.cn/#/exposome.

## Code availability

All the source code of the ExposomeX project is deployed on GitHub R package “ExposomeX” (see https://github.com/ExposomeX/Data-and-Code-for-ExposomeX-Manuscript.git). The source code works on Windows, macOS X, and most Linux distributions. The R code of the case study in R markdown format and the web module operation tutorials in PPT format were all provided at the website: http://www.exposomex.cn/#/toturial.

## Acknowledgements

We would like to express our gratitude to the working group of environmental exposure and human health of the China Cohort Consortium (http://chinacohort.bjmu.edu.cn/). Special thanks to the graduates (Mingyu Li, Baiqiang Li, Yizhong Jin, and Siqi Zhang from North China Electric Power University) for their hard work to develop the web-interface. We also thank Feng Zhao, Siyi Wang, and Jing Yang from Fudan University and Dr. Min Liu, Haoduo Zhao, Junjie Yang and Huili Du from Nanyang Technological University for participating in the work to establish the database and method development. Dr. Bin Wang was supported by the National Natural Science Foundation of China (Grant No. 42077390, 41771527) and Dr. Mingliang Fang was supported by the start-up grant from Fudan University (JIH1829010Y).

## Author contributions

B.W. and M.L.F. conceived the method and supervised its implementation. C.L., G.H.Z., M.Y. R., Y.Q.F., N.G., W.N. L., B.J., S.S., Y.T.W., H.W., F.R.Z., B.P., X.T.J., X.J.C., and N.M. developed the methods, packages, curated the databases and wrote the help documents and tutorials. C.L., Z.J.L., and X.Q.S built the websites. B.W. and M.L.F. prepared the figures and wrote the manuscript. All authors contributed to the final manuscript.

## Additional information

Supplementary information The online version contains supplementary material available at https://doi.org/XXXXXX. Correspondence and requests for materials should be addressed to Dr. Mingliang Fang.

