## Supplementary material for "ExposomeX: Integrative Exposomic Platform Expediates Discovery of “Exposure-Biology-Disease” Nexus": code

Supplementary Code_Fig3A-G_Stat

Supplementary Code_Fig3H-K_Viz

Supplementary Code_Fig4B_Cros

Supplementary Code_Fig4C_Surv

Supplementary Code_Fig4D_Panel

Supplementary Code_Fig4EF_NTA

Supplementary Code_Fig4G_Cros

Supplementary Code_Fig4H_MO

Supplementary Code_Fig4I_Mix

Supplementary Code_Fig4J_Medt

Supplementary Code_Fig5ABC_StatLink

Supplementary Code_Fig5DEF_Meta

Supplementary Code_Fig5GH_BioLink

Supplementary Code_Fig6ABC_Medt

Supplementary Code_Fig6D_Mix

Supplementary Code_Fig6E_BioLink

Supplementary Code_Fig6F_StatLink

Supplementary Code_Fig6G_Meta

Supplementary Code_Fig7A_MO

Supplementary Code_Fig7B_Viz

Supplementary Code_Fig7C_StatLink

Supplementary Code_Fig7D_Medt

Supplementary Code_Fig7E_Mix

Supplementary Code_Fig7F_NTA

*All the above files can be downloaded at the website:

<https://github.com/ExposomeX/Data-and-Code-for-ExposomeX-Manuscript.git>
