## Supplementary material for "ExposomeX: Integrative Exposomic Platform Expediates Discovery of “Exposure-Biology-Disease” Nexus": data

Supplementary Data_Fig3A-G_Stat

Supplementary Data_Fig3H-K_Viz

Supplementary Data_Fig4B_Cros

Supplementary Data_Fig4C_Surv

Supplementary Data_Fig4D_Panel

Supplementary Data_Fig4EF_NTA

Supplementary Data_Fig4G_Cros

Supplementary Data_Fig4H_MO

Supplementary Data_Fig4I_Mix

Supplementary Data_Fig4J_Medt

Supplementary Data_Fig5ABC_StatLink

Supplementary Data_Fig5DEF_Meta

Supplementary Data_Fig5GH_BioLink

Supplementary Data_Fig6ABC_Medt

Supplementary Data_Fig6D_Mix

Supplementary Data_Fig6E_BioLink

Supplementary Data_Fig6F_StatLink

Supplementary Data_Fig6G_Meta

Supplementary Data_Fig7A_MO

Supplementary Data_Fig7B_Viz

Supplementary Data_Fig7C_StatLink

Supplementary Data_Fig7D_Medt

Supplementary Data_Fig7E_Mix

Supplementary Data_Fig7F_NTA

*All the above files can be downloaded at the website:

<https://github.com/ExposomeX/Data-and-Code-for-ExposomeX-Manuscript.git>
