## Supplementary material for "ExposomeX: Integrative Exposomic Platform Expediates Discovery of “Exposure-Biology-Disease” Nexus": Tutoriols

- **Tutorials**

(1) Web modules

- Description: describe the data
- Visualization: visualize the data
- Database: support the ExposomeX platform
- Cross-sectional data: analyze the cross-sectional data
- Multi-omics data: analyze the data of multi-omics
- Nontarget data: analyze the non-targeted from HRMS
- Panel data: analyze the data of panel study
- Survival data: analyze the censored data
- Mediation effect: screening the mediation pathway
- Mixture effect: analyzing the effect of mixture exposure
- Statistical: explore the statistical explanation
- Biological: explore the biological explanation
- Meta-analysis: provide the summary of the previous publications

(2) R packages

- ExpoStatistics: describe the data
- ExpoViz: visualize the data
- ExpoDB: support the ExposomeX platform
- ExpoCros: analyze the cross-sectional data
- ExpoMultiomics: analyze the data of multi-omics
- ExpoNontarget: analyze the non-targeted from HRMS
- ExpoPanel: analyze the data of panel study
- ExpoSurvival: analyze the censored data
- ExpoMediation: screening the mediation pathway
- ExpoMixEffect: analyzing the effect of mixture exposure
- ExpoStatLink: explore the statistical explanation
- ExpoBioLink: explore the biological explanation
- ExpoMeta: provide the summary of the previous publications

* All the above up-to-date tutorial files can be downloaded at the website:

<http://www.exposomex.cn/#/toturial>
