## Supplementary material for "ExposomeX: Integrative Exposomic Platform Expediates Discovery of “Exposure-Biology-Disease” Nexus": Table S1

**Table S1**. Typical characteristics of the databases used for ExposomeX platform

| Classes ^a^ | Count | Descriptions of main sources ^b^ |
| --- | --- | --- |
| Exposure | 94,355 | STITCH, CTD, ToxCast, and T3DB. The chemical information are mainly from Chemical Translation Service (CTS). |
| Metabolite | 18,905 | KEGG for the human species. |
| Protein | 167,654 | UniprotKB in Uniprot consortium, STRING, and STITCH, and CTD for the human species. |
| Enzyme | 5,223 | KEGG for the human species. |
| Disease | 16,400 | Uniprot consortium, CTD and OMIM. |
| Chemical-protein | 2,053,899 | TTD, T3DB, CTD, ToxCast, and STITCH. |
| Protein-protein | 59,714 | STRING for the human species. |
| Protein-disease | 4,554 | Uniprot consortium for the human species |
| Chemical-GO | 25,096 | CTD |
| GO-Disease | 2,779,411 | CTD |
| Meta-analysis publications | 19,781 | CTD and Exposome-Explorer |

^a^ The related databases can be downloaded from the ExposomeX platform at the website:

<http://www.exposomex.cn/#/download>;

^b^ The related databases can be obtained through the following websites:

[HMDB: Human Metabolome Database (https://hmdb.ca/)](HMDB:%20Human%20Metabolome%20Database%20(https://hmdb.ca/))

[OMIM: Online Mendelian Inheritance in Man (https://omim.org/)](OMIM:%20Online%20Mendelian%20Inheritance%20in%20Man%20(https://omim.org/))

[STICH: Chemical-Protein Interaction Networks (http://stitch.embl.de/)](STICH:%20Chemical-Protein%20Interaction%20Networks%20(http://stitch.embl.de/))

UniProt consortium (https://www.uniprot.org/)

[STRING: Functional Protein Association Networks (https://cn.string-db.org/)](STRING:%20Functional%20Protein%20Association%20Networks%20(https://cn.string-db.org/))

[CTD: Comparative Toxicogenomics Database (http://ctdbase.org/)](CTD:%20Comparative%20Toxicogenomics%20Database%20(http://ctdbase.org/))

[T3DB: Toxin and Toxin Target Database (http://www.t3db.ca/)](T3DB:%20Toxin%20and%20Toxin%20Target%20Database%20(http://www.t3db.ca/))

[GO: Geneontology (http://geneontology.org/)](GO:%20Geneontology%20(http://geneontology.org/))

[KEGG: Kyoto Encyclopedia of Genes and Genomes (https://www.genome.jp/kegg/)](KEGG:%20Kyoto%20Encyclopedia%20of%20Genes%20and%20Genomes%20(https://www.genome.jp/kegg/))

[CTS: Chemical Translation Service (http://cts.fiehnlab.ucdavis.edu/)](CTS:%20Chemical%20Translation%20Service%20(http://cts.fiehnlab.ucdavis.edu/))

[ToxCast: Toxicity ForeCaster (https://www.epa.gov/chemical-research)](ToxCast:%20Toxicity%20ForeCaster%20(https://www.epa.gov/chemical-research))

TTD: Therapeutic Target Database (<http://db.idrblab.net/ttd/>)

Exposome-Explorer: (<http://exposome-explorer.iarc.fr/>)
