## Supplementary material for "ExposomeX: Integrative Exposomic Platform Expediates Discovery of “Exposure-Biology-Disease” Nexus": Figure S1

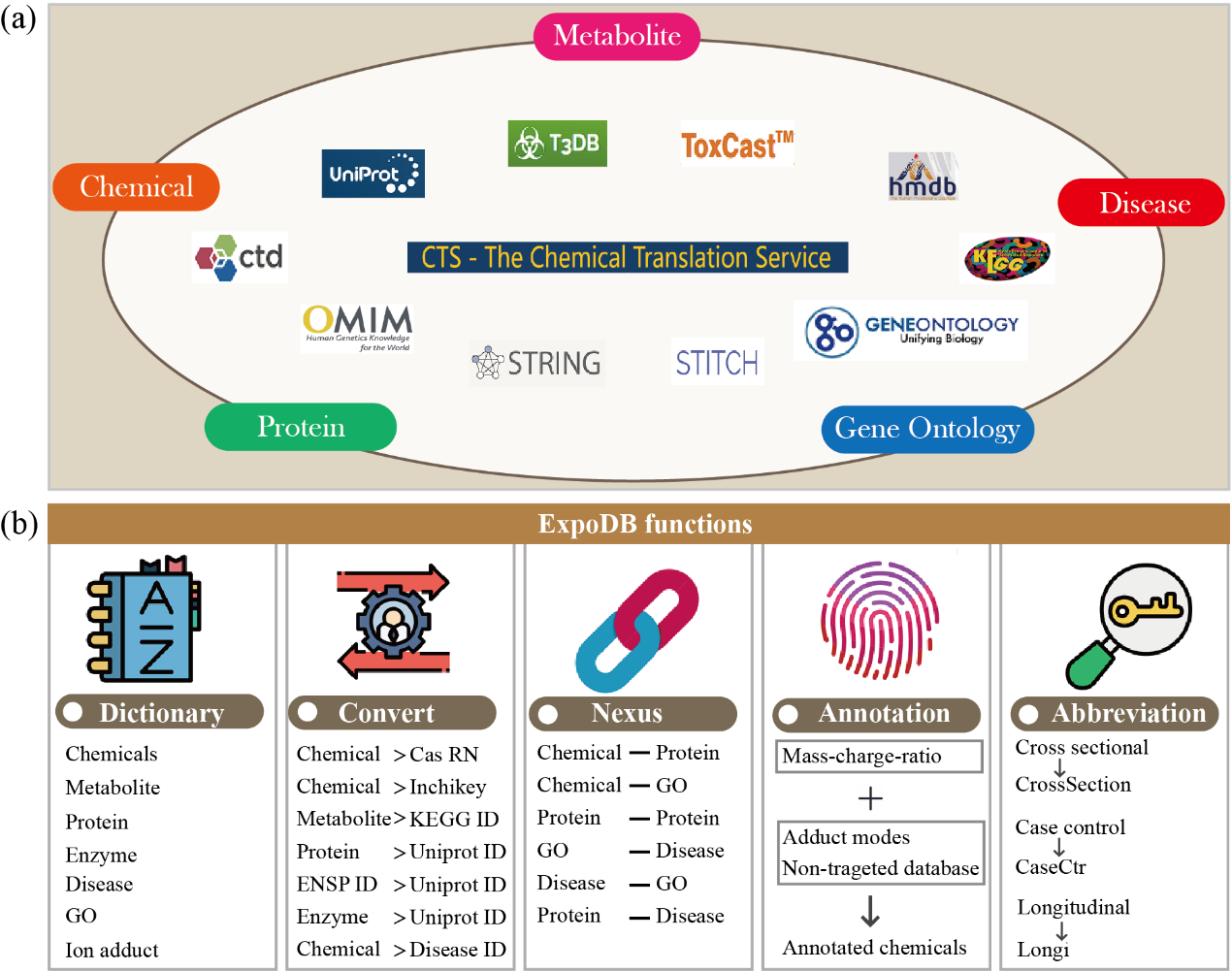

**Figure S1**. Databases used for ExposomeX platform development and five functions of ExpoDB module. (a) The databases used to develop ExposomeX platform; (b) The five functions of ExpoDB module. Of them, “Dictionary” is used for searching the related information about the databases, “Convert” for converting the names or IDs between different databases or nomenclatures, “Nexus” for exploring the potential nexuses between exposure, protein, phenotype, and disease, “Annotation” for annotating the non-target features from high-resolution mass spectrometry, “Abbreviation” for learning the nomenclature of "ExposomeX" platform.
